# Distilling Direct Effects via Conditional Differential Gene Expression Analysis

**DOI:** 10.1101/2025.09.29.678272

**Authors:** Jiaqi Gu, Andrew Skelton, James Staley, Pierce Popson, Lei Peng, Xiaoyu Song, Juliet Knowles, Zihuai He

## Abstract

Differential gene expression (DGE) analysis is foundational for interpreting RNA sequencing data, but it conflates direct biological effects with correlations propagated through gene co-expression. Across three RNA sequencing datasets (including a genome-scale perturb-seq experiment), we find that only a small fraction of differentially expressed genes have direct effects on the trait of interest, while the majority are undirected or passengers whose associations are mediated through other genes. To distinguish direct effect genes, we introduce conditional differential gene expression (CDGE) analysis, a framework that tests for conditional rather than marginal association between each gene and the trait of interest. Implemented via the GhostKnockoff procedure with lasso regression, CDGE delivers false discovery rate control, operates on summary statistics from existing DGE pipelines, and accommodates batch effects. The genes identified by CDGE mediate the effects of most other differentially expressed genes and show stronger enrichment for known protein-protein interactions and biological pathways than DGE-identified genes. These results suggest that the field has been systematically over-interpreting DGE outputs, and that distinguishing direct from mediated effects is essential for prioritizing genes for functional follow-up and therapeutic development.

## Introduction

The rapid development of RNA-sequencing (RNA-seq) technologies has provided extensive data on diseases and biological processes (Stark et al. 2019). RNA-seq is used to measure transcriptome expression levels in individual cells, enabling the study of cellular processes and biological pathways. Correspondingly, multiple statistical methods have been proposed to analyze these data, among which differential gene expression (DGE) analysis is one popular category (Kebschull et al., 2017; Costa-Silva et al., 2017; Jones et al., 2024; Rosati et al., 2024). DGE analysis identifies genes associated with traits of interest (e.g. diseases or gene perturbations) by comparing gene expression levels between cells of different conditions (e.g. healthy vs disease cells/tissues, control cells vs perturbed cells) under either parametric model (Ho et al., 2008; Robinson et al., 2010; Anders and Huber, 2010; Wang et al., 2010; Hardcastle and Kelly, 2010; Leng et al., 2013; Love et al., 2014; Ritchie et al., 2015; Ran and Daye, 2017) or nonparametric model (Tarazona et al., 2011; Li and Tibshirani, 2013; Tarazona et al., 2015).

These technologies have offered unprecedented opportunities to investigate and understand the underlying biological mechanisms of diseases or patterns of gene-gene interactions, especially with the aid of external information from functional enrichment analysis (Udhaya Kumar et al., 2020; Rosati et al., 2024).

Although widely utilized, most DGE analyses encounter difficulties in distinguishing between genes on the biological pathways and those not. This limitation arises because differentially expressed genes are identified based on their expression levels being marginally associated with the trait of interest, while the potential confounding brought by gene coexpression is ignored (Farahbod and Pavlidis, 2019). Instead of acting alone, genes and their corresponding proteins coordinate as functional modules or aggregate to form nano-machineries in biological processes (Lui et al., 2015). Thus, genes in same biological processes have similar expression patterns and strongly correlated expression levels (Boutros and Okey, 2005; Do and Choi, 2008). Without accounting for expression level correlations induced by functional interdependence, it is likely that genes not on the biological pathways are also identified as differentially expressed. For example, in Figure 1 (a) where we visualize gene-disease relations, there are 4 types of genes as follows.

- **Direct effect genes** are involved in the pathways of the disease and are the closest precedents of the disease (genes *x*_7_, *x*_8_,…,*x*_106_).
- **Indirect effect genes** are involved in the pathways of the disease and are not the closest precedents of the disease (genes *x*_1_, *x*_2_, …,*x*_6_).
- **Passenger genes** are not involved in the pathways of the disease but are affected by indirect effect genes or direct effect genes (e.g., genes *x*_107_, *x*_108_, …,*x*_306_).
- **No effect genes** are not involved in the pathways of the disease and are not affected by indirect effect genes or direct effect genes (genes *x*_307,_ *x*_308_,…,*x*_500_).

**Figure 1.**
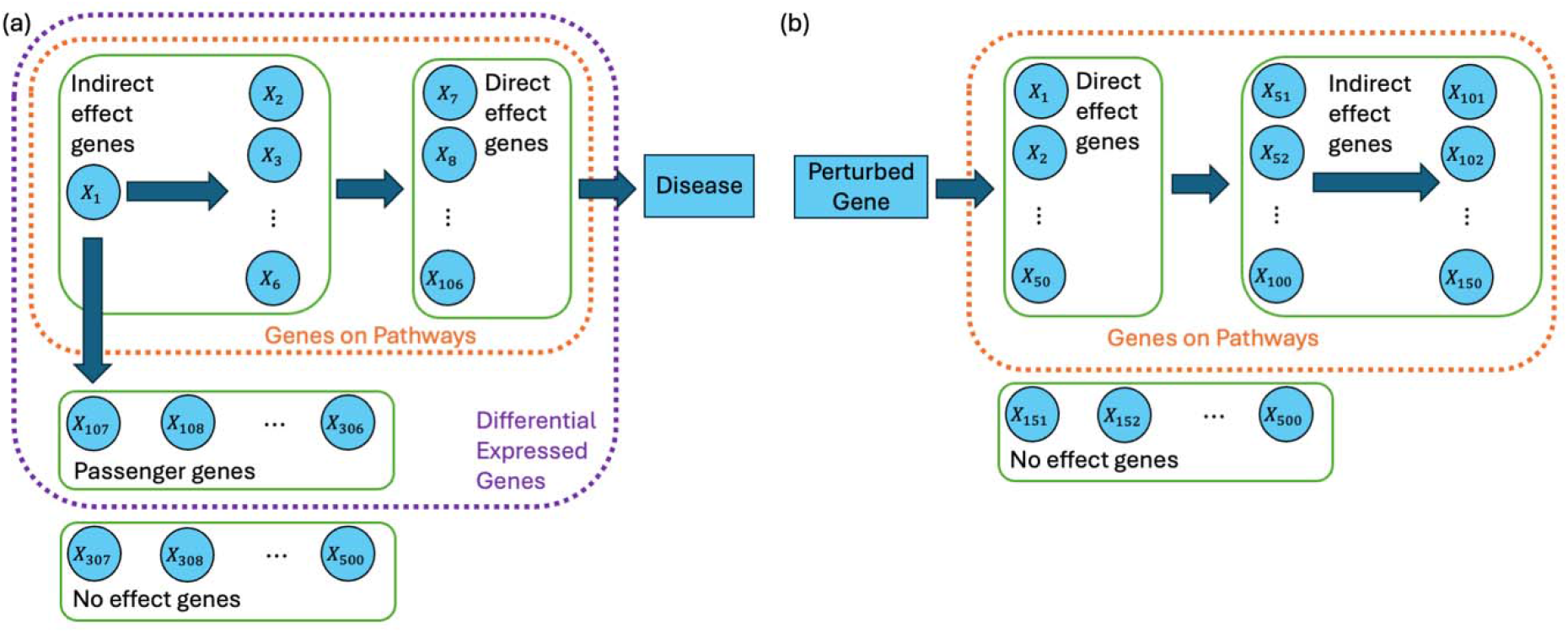
Graphical illustration of different types of genes when (a) investigating gene-disease relations or (b) investigating gene-gene relations.

Expression levels of the first three types of genes are expected to correlate with the disease of interest. We use the term “passenger” here in deliberate analogy to its usage in cancer genomics, where passenger mutations are correlated with the tumor phenotype but not causally upstream of it (Stratton et al., 2009; Vogelstein et al., 2013); we extend the term from the mutation context to the expression context, with the same intuition that a passenger is correlated with the trait of interest but downstream of it or genes on the pathways. As a result, the DGE analysis would identify not only the genes on the pathways (first two types) but also passenger genes (the third type). These redundantly identified genes would obscure the most important target genes when designing biological studies for functional validation or developing medicines. Under particular conditions, direct effects of genes can be translated to causal effects, and identifying genes with direct effects helps reveal the “wiring” of biological circuits and benefit medicine development. For example, therapies targeting direct effect genes or their products improve efficacy (Jiang et al., 2019) and reduce toxicity (Xu et al., 2022), with consistent benefits across different populations (Slamon et al., 2001; Slamon et al., 2011). In contrast, drugs targeting indirect effect genes or passenger genes show less consistent effects (Singh & Chakrabarti, 2023; Arias-Badia et al., 2025) and higher risks of adverse events (Donath & Shoelson, 2011). The need to identify direct effect genes also arises when we investigate gene expression level variations in genetic perturbation experiments (as visualized in Figure 1 (b)). Identifying direct effects from perturbed genes to measured genes can help to establish our knowledge of gene-gene relations and benefit medical practice. For example, in therapies that modify gene expression, we can monitor expression levels of only direct effect genes and truncate unwanted mutation effects with fewer drugs. This calls for statistical approaches that can pinpoint genes with direct effects.

In this article, we argue that DGE analysis identifies the right set of trait-relevant genes but cannot distinguish drivers from passengers within that set, leading to systematic misallocation of follow-up effort, from gene prioritization to drug-target selection. Beyond the methodological contribution, the four-category taxonomy we introduce (direct, indirect, passenger, no-effect genes) provides a vocabulary for reasoning about what differential expression results actually mean biologically. We then present conditional differential gene expression (CDGE) analysis as a framework for separating direct from mediated and passenger effects. CDGE tests the conditional association between each gene and the trait of interest, given all other genes, and we discuss the causal interpretation of these conditional effects under stated assumptions.

To implement CDGE, we adapt the recently developed GhostKnockoff procedure with lasso regression (Chen et al., 2024; He et al., 2024). Our approach takes Z-scores from existing DGE pipelines and the gene expression correlation matrix as inputs, produces knockoff copies of the Z-scores as synthetic controls, and applies the knockoff filter (Candès et al., 2018) to identify direct effect genes with finite-sample false discovery rate control. Because inference operates on summary statistics, the method applies to datasets with cell-level records as well as to datasets where only summary statistics are available, and it accommodates batch effects through the choice of upstream DGE approach. We validate CDGE in simulations and across three RNA sequencing datasets of increasing scale. The headline application is a genome-scale perturb-seq experiment (Replogle et al., 2022) covering 1,284 perturbations: CDGE reduces the number of identified gene-gene connections by approximately 70%, and the retained connections explain over 75% of the connections identified by standard DGE while showing markedly higher enrichment for known protein-protein interactions. Two scRNA-seq applications, one in human T-cells from psoriasis patients and one in a Scn8a+/-mouse model of epilepsy, reproduce this pattern: CDGE-identified genes mediate the effects of the large majority of other differentially expressed genes and concentrate biological signal in pathways relevant to disease.

## Results

We validate CDGE in four settings of increasing scale and decreasing access to ground truth. We first describe the framework and its causal interpretation, then evaluate it in a controlled simulation that establishes finite-sample false discovery rate control. We next apply CDGE to two single-cell disease studies, one in human T-cells from psoriasis patients and one in a Scn8a+/-mouse model of epilepsy, that test the method on observational human and mouse data with strong prior biological knowledge. Finally, we apply CDGE to a genome-scale perturb-seq experiment with 1,284 perturbations (Replogle et al., 2022), which tests its ability to identify direct gene-gene regulatory effects across thousands of designed interventions and provides the most stringent external validation against the STRING protein-protein interaction network.

### Conditional differential gene expression (CDGE) analysis via Ghostknockoff

Usual DGE analysis infers marginal correlations between the trait of interests (denoted by *Y*) and the expression level of different genes (denoted by *x*_*j*_’s) via testing *H*_*M,j*_: *X*_*j*_ ⊥ *Y, j* = 1,…,*p*. In this article, we propose the CDGE analysis that investigates the conditional association between each gene and the trait of interests conditioning on all other genes by testing

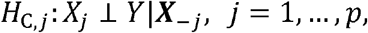

where ***X***_-*j*_ encodes expression levels of all but the *j*-th genes. That is to say, CDGE analysis aims to identify whether (the distribution of) *Y* would change when the target gene up-regulates or down-regulates given expression levels of all other genes remain unchanged. As shown in Figure 1 (a), when expression levels of all other genes remain unchanged, regulation of the target gene would impact *Y* only via its direct effect if exists. Thus, *H*_C,*j*_depicts the existence of the *j*-th gene’s direct effect (if *H*_c,*j*_ is true, the *j*-th gene has no direct effect; if *H*_C,*j*_is false, the *j*-th gene has direct effect) and CDGE analysis can distinguish these direct effects from effects mediated via other genes, providing more information of the underlying biological mechanism. The interpretation of *H*_C,*j*_is similar when we investigate gene-gene connections in Figure 1 (b), except that regulation of the measured gene *X*_*j*_ would be impacted by *Y* (indicating whether the perturbation target gene is perturbed or not) if *H*_C,*j*_is false and direct effect from the perturbation target gene to the measured gene exists.

In this article, we perform knockoff-based CDGE analysis by adopting the recently developed Ghostknockoff with lasso regression (Chen et al., 2024; He et al., 2024). Overview of our knockoff-based CDGE analysis is provided in Figure 2, while details and theoretical reasoning of its FDR control with respect to *H*_c,1_, …,*H*_c,*p*_ are provided in the “Methods” section. In the first step, we perform existing DGE analysis and compute p-values of differential expression for all genes (*pval*, _1_,…,*pval*, _*p*_). We then transfer all p-values into Z-scores via the inverse cumulative distribution function of standard normal distribution (Φ^-1^ (·)) as

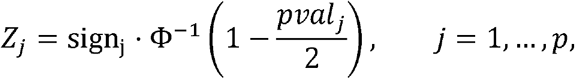

where sign_*j*_ depicts whether the *j*-th gene in cells of contrasted condition is upregulated or downregulated compared to cells of baseline condition. At the same time, the shrinkage estimation of correlation matrix (denoted by **Σ)** is computed (Ledoit and Wolf, 2003; Schäfer and Strimmer, 2005) using expression levels of all genes in all cells. With Z-score vector **z** = (*Z*1,…,*Z*_*p*)_^T^ and the estimated correlation matrix **Σ** as inputs, the knockoff-based CDGE analysis

1. generates knockoff copy 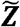 that serves as synthetic control;
2. fits a lasso regression model using summary statistics and obtain coefficient estimates, which empirically depict direct effects of genes and their knockoff copies to the trait of interest;
3. applies the knockoff filter to select direct effect genes under the target FDR level.

**Figure 2.**
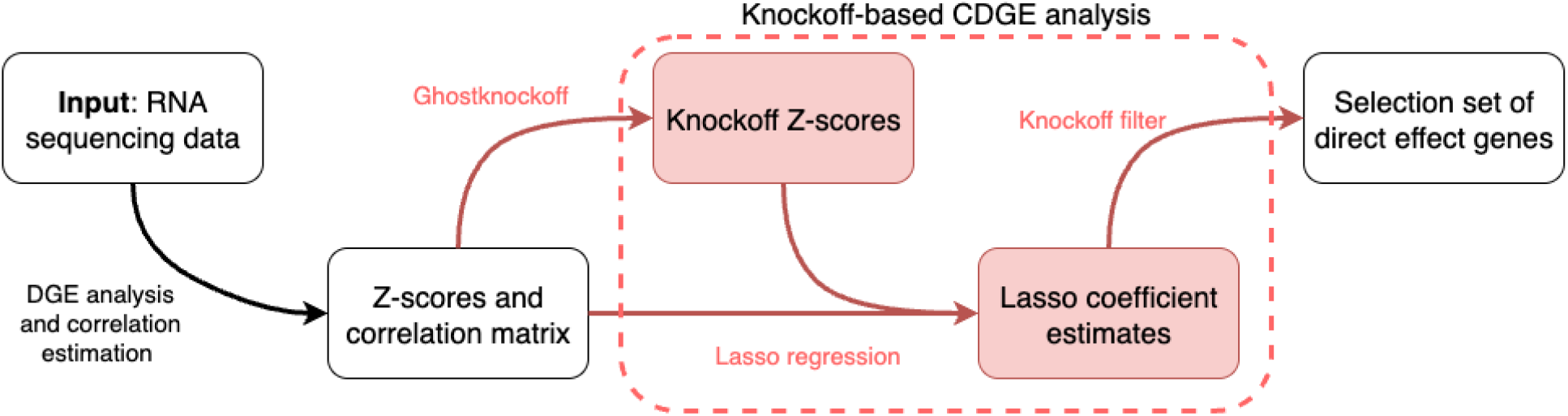
Overview of the knockoff-based CDGE analysis.

It is worth noting that our knockoff-based CDGE analysis encodes all data information into Z-scores and the correlation matrix of expression levels for inference. This property gives our approach the flexibility to be applied to either datasets with cell-level records (as in the psoriasis and epilepsy applications) or with only summary statistics available (as in the perturb-seq application). As shown in Chen et al. (2024), as long as the correlation matrix is well-estimated, the generated knockoff copy satisfies the extended exchangeability property (Yu et al., 2024) and the obtained set of direct effect genes by the knockoff-based CDGE analysis is equivalent to the second order knockoffs (Candès et al., 2018) which needs access of cell-level records. Thus, our knockoff-based CDGE analysis possesses valid FDR control with respect to ‘s without substantial power loss. Another benefit of using Z-scores to encode information is that we can incorporate existing DGE approach to address any possible batch effects in single cell datasets. By doing so, we do not need to care about batch effects in the generation of knockoff copy. In contrast, batch effects bring dependency among records of different cells and complicate the generation of cell-level knockoff copies of gene expression levels. This separation of concerns is a structural advantage of the summary-statistics formulation: the upstream DGE model handles cell-level dependencies and confounders, while CDGE operates downstream on Z-scores. Any batch-handling capability of the chosen DGE method (e.g., the design-matrix adjustment in DESeq2, mixed-effects models for repeated measures, or factor-model approaches for unmeasured confounders) is automatically inherited by CDGE, without requiring re-derivation of the knockoff construction for each new design.

### Causal interpretation of CDGE analysis

In general, the conditional association between genes and the trait depicted by false cannot be simply interpreted as causality, unless further assumptions are imposed. Under the potential outcomes framework (Rubin, 1974; Rubin, 2005; Imbens and Rubin, 2015), we can let denote the potential binary outcome of a cell that would have been observed if the gene expression levels are. Under the assumptions that

1. (**Absence of unmeasured confoundings**) ;
2. (**Positivity**) for all possible values of ;
3. (**Consistency**) the observed if, the conditional association between gene *x*_*j*_ and *Y* can be translated to causal effects that

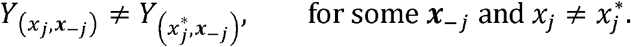

We emphasize that these assumptions, in particular the absence of unmeasured confounding, are unlikely to hold exactly in observational transcriptomics. Unmeasured cell-state variables, technical batch factors, and shared upstream regulators can all violate this assumption, and positivity is similarly difficult to verify when expression levels are continuous and high-dimensional. We therefore view the causal interpretation as an idealized benchmark rather than a routine claim of CDGE outputs. In practice, the conditional independence tested by CDGE should be understood as identifying direct statistical effects under the assumed correlation structure, with perturb-seq experiments providing the closest realistic approximation to the causal interpretation because the perturbation itself is a designed intervention. For purely observational scRNA-seq applications, conditional effects are best read as more biologically informative than marginal ones, without claiming full causal identification.

### Simulation Verification that CDGE analysis Identifies Direct Effects

To validate our statement that CDGE analysis can identify direct effect genes, we conduct a simulated experiment with *p* = 500 genes and *n* = 5,000 cells. Specifically, expression levels of these *p* = 500 genes are generated from the hierarchical model as visualized in Figure 3 (a) and detailed in “Experiments” section, while the response *Y* is generated from the logistic model that

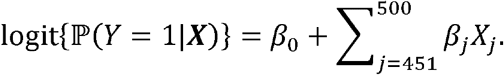

**Figure 3.**
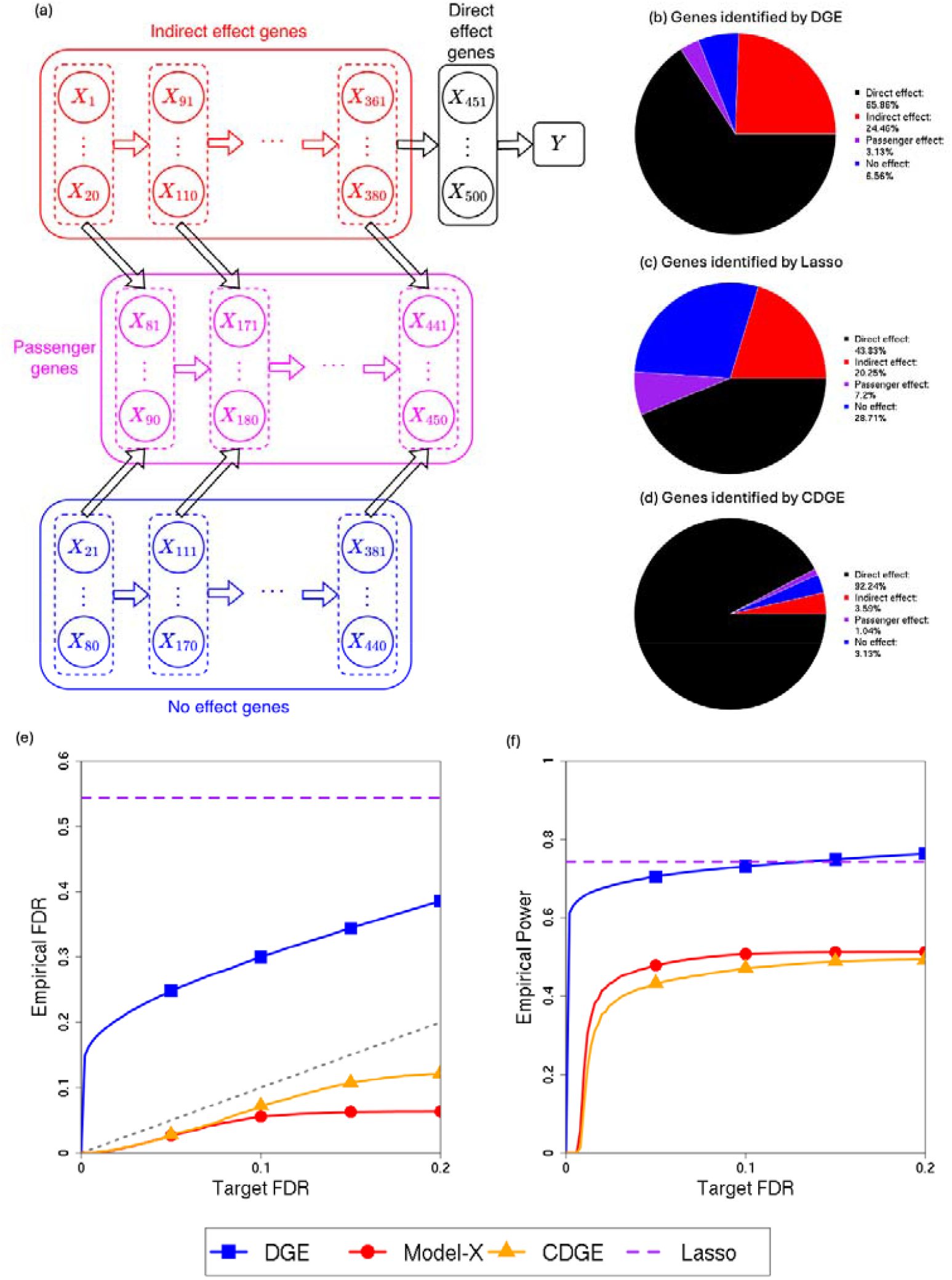
**(a)** Graphical illustration of the hierarchical model under which expression levels of *p* = 500 genes and the response are generated. **(b)** Average proportions of different types of DGE-significant genes under the FDR level 0.10 with BH correction. **(c)** Average proportions of different types of genes identified as direct effect genes under penalize regression. **(d)** Average proportions of different types of CDGE-identified genes under the FDR level *α* = 0.10 with the knockoff-based procedure. **(e)** Empirical FDR of DGE, penalized regression, model-X knockoffs and knockoff-based CDGE analysis in identifying direct effect genes over all target FDR levels *α* ∈[0,0.2]. **(f)** Empirical power of DGE, penalized regression, model-X knockoffs and knockoff-based CDGE analysis in identifying direct effect genes over all target FDR levels *α* ∈[0,0.2].

Here, *β*_451_, …,*β*_500_ are i.i.d. samples from the normal distribution N(0,0.02) and *Y* denotes an indicator of whether the cell comes from the disease group. By doing so, we simulate all four types of genes with respect to *Y* (direct effect genes, indirect effect genes, passenger genes and no effect genes). For example, *x*_451_,…,*x*_500_ correspond to direct effect genes and x_361_,…,x_380_ are indirect effect genes which affect both direct effect genes and passenger genes *x*_441_,…,*x*_450_. To mimic the complexity of biological pathways, we set 5 layers of indirect effect genes. While existing DGE tests are anticipated to identify direct effect genes, indirect effect genes and passenger genes (respectively colored black, red and purple in Figure 3 (a)), we expect CDGE analysis can narrow down the focus to identify direct effect genes (*X*_451_, …,*X*_500_). We perform DGE analysis on this data to compare gene expression levels between cells in the disease group (*Y* = 1) and the control group (*Y* = 0) via the Wilcoxon rank-sum test and we compute the Z-score of the *j*-th gene as *z*_*j*_ for *j*= 1,…,*p*.

Following the above procedure, we generate 1,000 replicates. We first display in Figure 3 (b) the average proportions of different types of DGE-significant genes under the FDR level 0.10 with the Benjamini-Hochberg (BH; Benjamini and Hochberg, 1995) correction. Although DGE analysis manages to identify a lot of genes on the pathways (direct effect genes and indirect effect genes), direct effect genes account for only 65.86% of DGE-identified genes. That is to say, DGE analysis cannot distinguish direct effect genes from other genes with differential expression (indirect effect genes and passenger genes account for 24.46% and 3.13% respectively). To account for correlations among expression levels, penalized regression is another popular category of approaches for direct effect genes identification. However, when we apply the logistic regression with lasso penalty, only 43.83% of identified genes are direct effect genes as shown in Figure 3 (c). In short, neither DGE analysis nor penalized regression provide FDR-controlled detection of direct effects.

We implement our knockoff-based CDGE analysis to identify direct effect genes at the same target FDR level *α* = 0.10. To validate that these CDGE-identified genes mainly correspond to those of direct effects, we present the average proportions of different types of genes in these CDGE-identified genes over 1,000 replicates in Figure 3 (d).We find that although both the knockoff-based CDGE analysis and DGE analysis only identify several no effect genes, CDGE analysis prioritizes direct effects. Specifically, over 1,000 replicates, the average proportion of CDGE-identified genes without direct effects is only 7.76% (while there are 90% genes without direct effects among all genes), suggesting that CDGE analysis does identify direct effect genes with valid FDR control. Such an FDR control remains valid throughout all target FDR levels *α* ∈ [0,0.2] as displayed in Figure 3 (e), in comparison to the inflated FDR of both DGE analysis and lasso regression.

Because our knockoff-based CDGE analysis is established upon the framework of Ghostknockoff with lasso regression (Chen et al., 2024; He et al., 2024) and is a summary-statistics method, we also compare it with the model-X knockoffs to investigate the power loss to apply a summary-statistics method when cell-level data are available. Specifically, in the implementation of model-X knockoffs, we adopt the second-order construction of Candès et al. (2018) in synthesizing knockoff copies of gene expression levels ***X*** and fit a logistic regression model with lasso penalty to obtain estimates 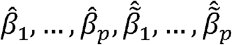 for inference. As shown in Figure 3 (f), we find that our knockoff-based CDGE analysis only suffer minor loss with respect to the model-X knockoffs. That is to say, although the knockoff-based CDGE analysis is implemented on the summary statistics ***Z***, it does not miss substantial information related to genes’ direct effect.

### Applications to RNA sequencing data

To illustrate that CDGE analysis does empirically uncover the biological mechanisms by prioritizing genes with direct effects, we perform knockoff-based CDGE analysis on three RNA sequencing datasets.

#### Psoriasis dataset

We consider the single-cell RNA-seq dataset collected by Reynolds et al. (2021) in the study of inflammatory skin diseases. With the focus on psoriasis, a chronic autoimmune disease leading to abnormal patches on skin, we extract gene expression records of 120,309 skin cells from three psoriasis patients (denoted as P1, P2 and P3). Note that for each patient, epidermis and dermis cells in both lesional and non-lesional regions are collected and sequenced. Expression levels of 28,383 genes are measured by counts of corresponding mRNA over all cells. Based on the knowledge that psoriasis is a well-understood homogeneous disease with highly successful therapeutics around the T-cell axis, we focus our analysis on 23,090 T-cells. The numbers of different types of T-cells from different patients are summarized in Figure 4 (a). For numerically stable inference and better biological relevance, we filter out genes that pass the Cook’s distance threshold or express in less than 1% cells, leading to 9,661 genes remaining in our analysis.

**Figure 4.**
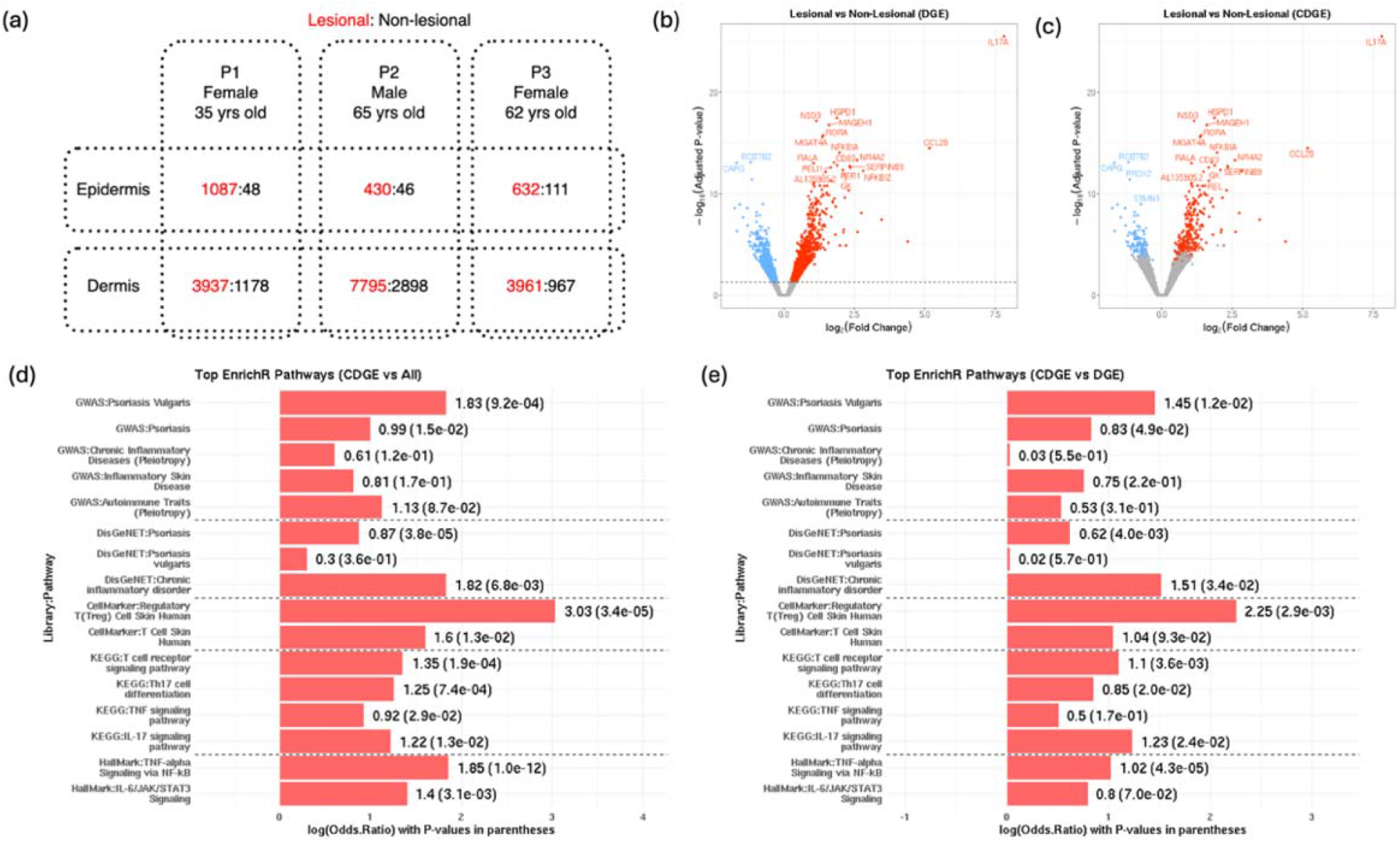
**(a)** The number of epidermis and dermis T-cells in both lesional and non-lesional regions from different patients. **(b)** Volcano plot of DGE analysis, where DGE-significant genes (under the target FDR level 0.05 with BH correction) are highlighted in red (upregulation) and blue (downregulation). **(c)** Volcano plot of knockoff-based CDGE analysis, where CDGE-significant genes (under the target FDR level *α* = 0.05) are highlighted. **(d)** Enrichment analysis of CDGE-identified genes in biological pathways related to psoriasis, inflammatory diseases, autoimmune traits and skin T-cells (compared to all genes). **(e)** Enrichment analysis of CDGE-identified genes in biological pathways related to psoriasis, inflammatory diseases, autoimmune traits and skin T-cells (compared to DGE-identified genes).

To begin with, we perform the pseudo-bulk DGE analysis using DESeq2 to investigate the marginal association between gene expression levels and the lesional state of cells, adjusting for their origins (P1, P2 or P3) and anatomic states (dermis or epidermis). We identify 2,509 differentially expressed genes under the target FDR level 0.05 with the BH correction. Volcano plot of the DGE analysis is provided in Figure 4 (b) and details of these differentially expressed genes are in Supplementary Table S1. As differentially expressed genes can be either with direct effects, with indirect effects or with passenger effects (shown in Figure 1 (a)), we can’t tell which genes have direct effects from Figure 4 (b). Genes with smallest p-values are possible to be those with indirect effects or passenger effects. For example, among the identified differentially expressed genes, IL17A, CCL20, IL23R and IL22 are known to be on biological pathways of psoriasis in an extensive literature, and the corresponding targeted gene therapies have been shown to be effective (Furue et al., 2020; Reynolds et al., 2021; Ghoreschi et al., 2021; Elnabawi et al., 2021). However, they may not be the top signals with smallest p-values in the DGE analysis, and it is difficult to determine whether their effects are direct effects. For example, IL23R and IL22 have the 77-th and 752-th smallest p-values respectively. We conduct the knockoff-based CDGE analysis on DGE Z-scores and expression level correlation matrix **Σ** of all genes to distill direct effects. Considering that cells from the same patients are not independent, we estimate the correlation matrix **Σ** as a factor model in the similar way of Leek and Storey (2007) and Fan et al (2020). Figure 4 (c) highlights all 378 CDGE-identified genes (under the target FDR level *α* = 0.05) in the volcano plot, whose details can also be found in Supplementary Table S1. Genes IL17A, CCL20 and IL23R are significant in the knockoff-based CDGE analysis, suggesting that they are more likely to have direct effects. To validate that these identified direct effect genes mediate the effects of the remaining differentially expressed genes, we adjust for CDGE-identified genes and perform DGE analysis of all other genes. After adjusting for 378 CDGE-identified genes, the remaining genes lose statistical significance in the DGE analysis, a pattern consistent with the 378 CDGE-identified genes mediating the 2,131 remaining differentially expressed genes’ effect on psoriasis. We note that conditioning on a large covariate set reduces statistical power generally, so the strongest causal-mediation interpretation requires further validation; the result we report should be read as evidence that CDGE-identified genes capture the bulk of the trait-relevant signal in this dataset, rather than as a definitive causal claim. Such a phenomenon is reproducible among different runs of knockoff-based CDGE analysis (which is random due to the knockoffs generation).

As the CDGE-identified genes empirically mediate all gene effects on psoriasis, we expect that these genes are more biologically informative with respect to psoriasis. To verify this, we perform enrichment analysis using multiple biological process pathways related to psoriasis, inflammatory diseases, autoimmune traits and skin T-cells. These pathways come from different libraries of the “Enrichr” database (Chen et al., 2013; Kuleshov et al., 2016; Xie et al., 2021). Figures 5 (d)-(e) present the enrichment of CDGE-identified genes over psoriasis-related pathways, in comparison against all genes and DGE-identified genes respectively. In general, CDGE-identified genes are enriched in all relevant pathways. For example, our CDGE identifies fundamental gene RNF114 for T-cell activation (Tsoi et al., 2012), gene IL23R for the functional receptor for cytokine IL-23 that drives the differentiation and maintenance of Th17 cells (Duerr et al., 2006) and gene REL for transcription factor of cytokines IL-12 and IL-23 (Gilmore and Gerondakis, 2011). In contrast, differentially expressed genes STAT3 and JAK1 used as drug targets are not identified in CDGE analysis, suggesting these genes are likely to have no direct effects to psoriasis.

**Figure 5.**
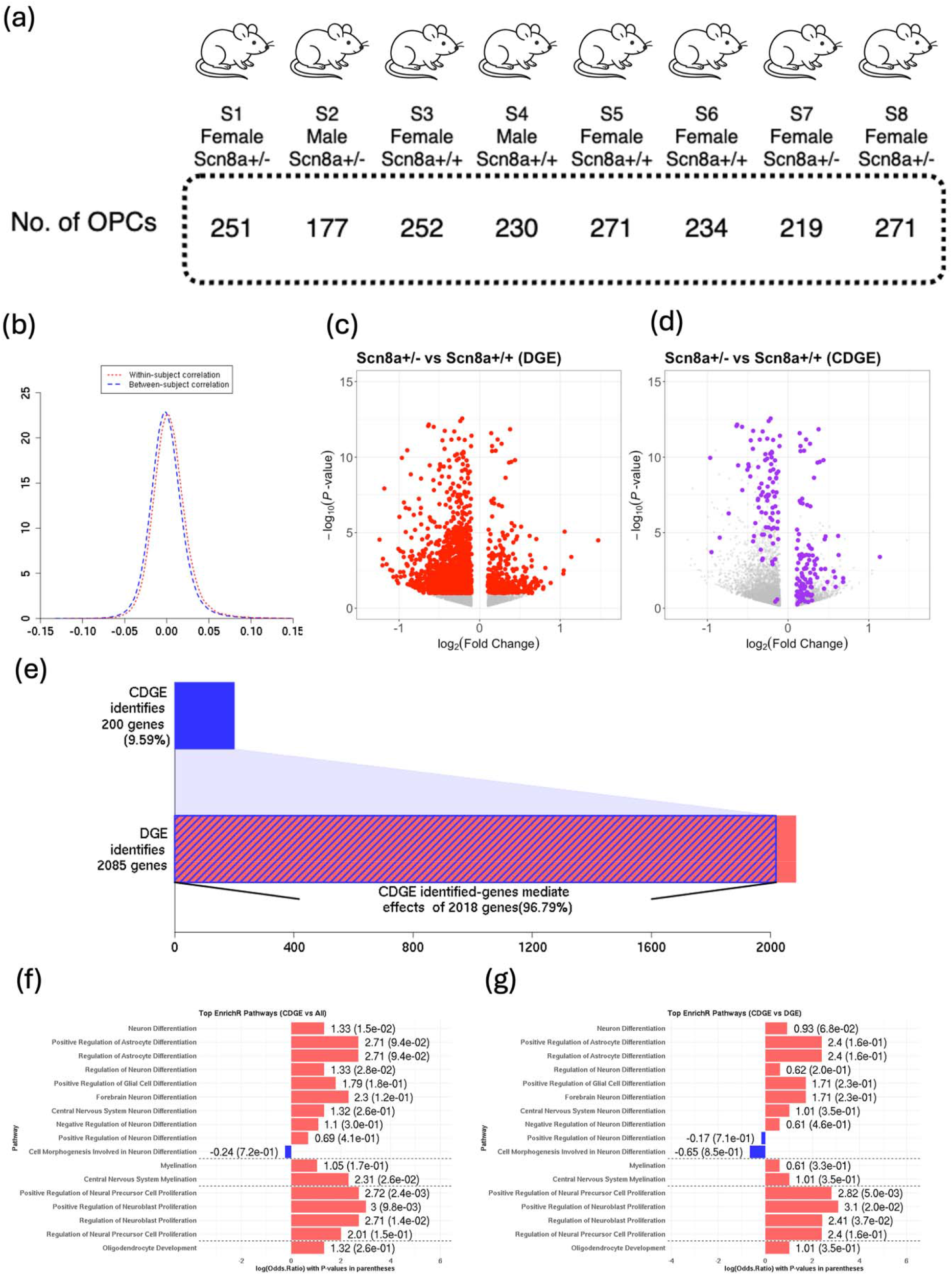
**(a)** Genotype, sex and the number of OPCs for each mouse. **(b)** Kernel density estimates of the distribution of gene expression correlation between cells from the same mouse (red dotted line) or between cells from different mice (blue dashed line). **(c)** Volcano plot of DGE analysis, where DGE-significant genes (under the target FDR level 0.10 with BH correction) are highlighted in red. **(d)** Volcano plot of knockoff-based CDGE analysis, where CDGE-significant genes (under the target FDR level *α* = 0.10) are highlighted in purple. **(e)** Comparison of the number of CDGE-identified genes, the number of DGE-identified genes and the number of genes whose effect is mediated by CDGE-identified genes. **(f)** Enrichment analysis of CDGE-identified genes in biological process pathways of neural cell differentiation, myelination, proliferation and oligodendrocyte development (compared to all genes). **(g)** Enrichment analysis of CDGE-identified genes in biological process pathways of neural cell differentiation, myelination, proliferation and oligodendrocyte development (compared to DGE-identified genes).

#### Epilepsy dataset

We also apply the knockoff-based CDGE analysis to a single-nucleus RNA-seq (snRNA-seq) dataset collected in investigation of biological mechanisms underlying epilepsy progression. Epilepsy is a type of non-communicable neurological disorder defined by the predisposition for recurrent, unprovoked seizures (Srivastava et al, 2021; Knowles et al., 2022). This snRNA-seq dataset is collected using a genetic mouse model of generalized epilepsy, *Scn8a*+/-. These mice express a loss of function mutation in *Scn8a*, which encodes the voltage-gated sodium channel alpha subunit (*Na*_v_ 1.6). Such a loss of function mutation leads to spontaneous, progressive absence seizures (Papale et al., 2009; Makinson et al., 2017; Knowles et al., 2022). To understand biological pathways of epilepsy progression in the *Scn8a*+/-model, we sequence expression levels of 57,010 genes over 63,141 cells from 8 mice, including 4 *Scn8a*+/-mice and 4 *Scn8a*+/+ littermate control mice. Details of our biological experiment are provided in the “Experiments” section. Expression levels of all genes are measured by counts of corresponding mRNA molecules over all cells. Motivated by the previous discovery (Knowles et al., 2022) that seizures in *Scn8a*+/-mice induce activity-dependent proliferation and maturation of oligodendrocyte precursor cells (OPCs), as well as increase callosal myelination which in turn, promotes further progression of epilepsy, we focus on differentially expressed genes in OPCs (1,905 in total). Genotypes, sex of these 8 mice and the number of their OPCs is summarized in Figure 5 (a).

For numerical stability of analysis, we filter out all genes with no more than 5% nonzero counts among the 1,905 OPCs. There remain 6,163 genes in our analysis. Before we perform the DGE analysis, we first investigate whether cells from the same mice are correlated or not. As shown in Figure 5 (b), gene expression correlation between cells from the same mouse follows the same distribution as the gene expression correlation between cells from different mice, suggesting that the within-subject dependency of gene expression levels is neglectable. Based on this, we perform the DGE analysis with univariate Poisson regression models for all genes. Under the target FDR level 0.10 with BH correction, 2,085 genes are identified as differentially expressed genes between heterozygous OPCs (from Scn8a+/-mice) and wild-type OPCs (from Scn8a+/+ mice), which are shown in Figure 5 (c) and detailed in Supplementary Table S2. These identified differentially expressed genes can be either direct effect genes, indirect effect genes or passenger effects. To identify direct effect genes, we apply knockoff-based CDGE analysis on DGE Z-scores and expression level correlation matrix **Σ** of all genes. Here, **Σ** is computed via the shrinkage estimation approach (the R package “corpcor”; Schäfer and Strimmer, 2005) using all cells. Figure 5 (d) highlights all 200 CDGE-identified genes (under the target FDR level *α* = 0.10) in the volcano plot, whose details are provided in Supplementary Table S2. Accounting for correlations of expression levels in our analysis, we eliminate genes’ marginal associations with epilepsy that are brought by their correlations to direct effect genes. As a result, we identify 200 direct effect genes (26 of which are not identified in DGE analysis) while 1,911 significantly differentially expressed genes are not identified in the knockoff-based CDGE analysis. This suggests that more than 90% of differentially expressed genes have either indirect effects or passenger effects mediated by CDGE-identified genes. To validate this statement, we adjust for 200 CDGE-identified genes and perform DGE analysis on the remaining genes again. As shown in Figure 5 (e), marginal associations between 96.79% of differentially expressed genes and epilepsy are decreased by adjusting for the 200 CDGE-identified genes. This validates our previous statement that effects of more than 90% differentially expressed genes are mediated.

As CDGE analysis targets direct effect genes, it is expected that CDGE-identified genes are more informative with respect to biological processes of OPCs, including proliferation, differentiation and myelination. To verify the biological relevance of CDGE-identified genes, we perform an enrichment analysis. Here, we use the gene set library “GO_Biological_Process_2025” (Ashburner et al., 2000; Aleksander et al., 2023) in the “Enrichr” database (Chen et al., 2013; Kuleshov et al., 2016; Xie et al., 2021) for analysis. Figures 5 (f)-(g) present the enrichment of CDGE-identified genes over pathways of neuron proliferation, neuron differentiation, myelination and oligodendrocyte development, in comparison against all genes and DGE-identified genes respectively. In general, CDGE-identified genes are enriched in almost all relevant pathways, especially in some differentiation pathways and all proliferation pathways, no matter compared against all genes or DGE-identified genes. Although not very large, log(OR) of myelination and oligodendrocyte development are positive with values around 1 when comparing CDGE-identified genes against DGE-identified ones. For example, our CDGE identifies fundamental genes for myelin sheaths formation and maintenance in deep white matter (MBP and PLP1) and early-stage differentiation of OPCs (NCAM1) (Harlow et al., 2014; Tisoncik-Go et al. 2024). All these results are consistent to the existing discovery that epilepsy causes hypermyelination of seizure circuits related to neuron differentiation, myelination and oligodendrocyte development. In contrast, differentially expressed genes SLC17A7 and CPLX1, which relate to oligodendrocyte lineage and synaptic signaling, are not identified in CDGE analysis, suggesting these genes are not likely to have direct effects to epilepsy. Such a case is true even for several down-regulated genes with the smallest p-value in the DGE test, which means that the most significant differentially expressed genes may fail to capture direct effects.

#### Application to genome-scale perturb-seq experiments

In addition to RNA sequencing datasets that investigate gene-disease relations, we apply the knockoff-based CDGE analysis to the genome-wide perturb-seq dataset collected by Replogle et al. (2022) that consists of 2,273 gene perturbation experiments for gene-gene connection study. To investigate functions of poorly characterized genes and uncover new regulators of various molecules activities (e.g., ribosome biogenesis, transcription and mitochondrial respiration), Replogle et al. (2022) collected expression levels of human genes within CRISPR interfered cells in chronic myeloid leukemia and retinal pigment epithelial cell lines. Specifically, for each gene of interest, they perform loss-of-function perturbation on it in several cells (5∼1,996) and compare the expression levels of all genes in these cells to control cells which are not perturbated. In our analysis, we use the pseudo-bulk dataset that records pseudo-bulk gene expressions in cells sampled in the sixth day after transduction. DGE Z-scores of 8,563 genes are provided in Replogle et al. (2022) by comparing the perturbed cells in each experiment to the 10,691 control cells via the *t*-test. We filter out experiments with potentially inflated Z-scores, defined as those with median of 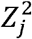 larger than 70%-quantile of 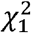 distribution. This results in 1,284 “P1P2” experiments with perturbation targeted at the consensus transcription start site, which would simultaneously perturb the primary and the secondary transcription start sites that are less than 1 kb apart (Horlbeck et al., 2016). Brief visualization of these Z-scores is provided in Figure 6 (a). Among the experiments, 123,043 gene-gene connections (95.83 per experiments) are identified by DGE analysis under the target FDR level 0.05 with BH correction.

**Figure 6.**
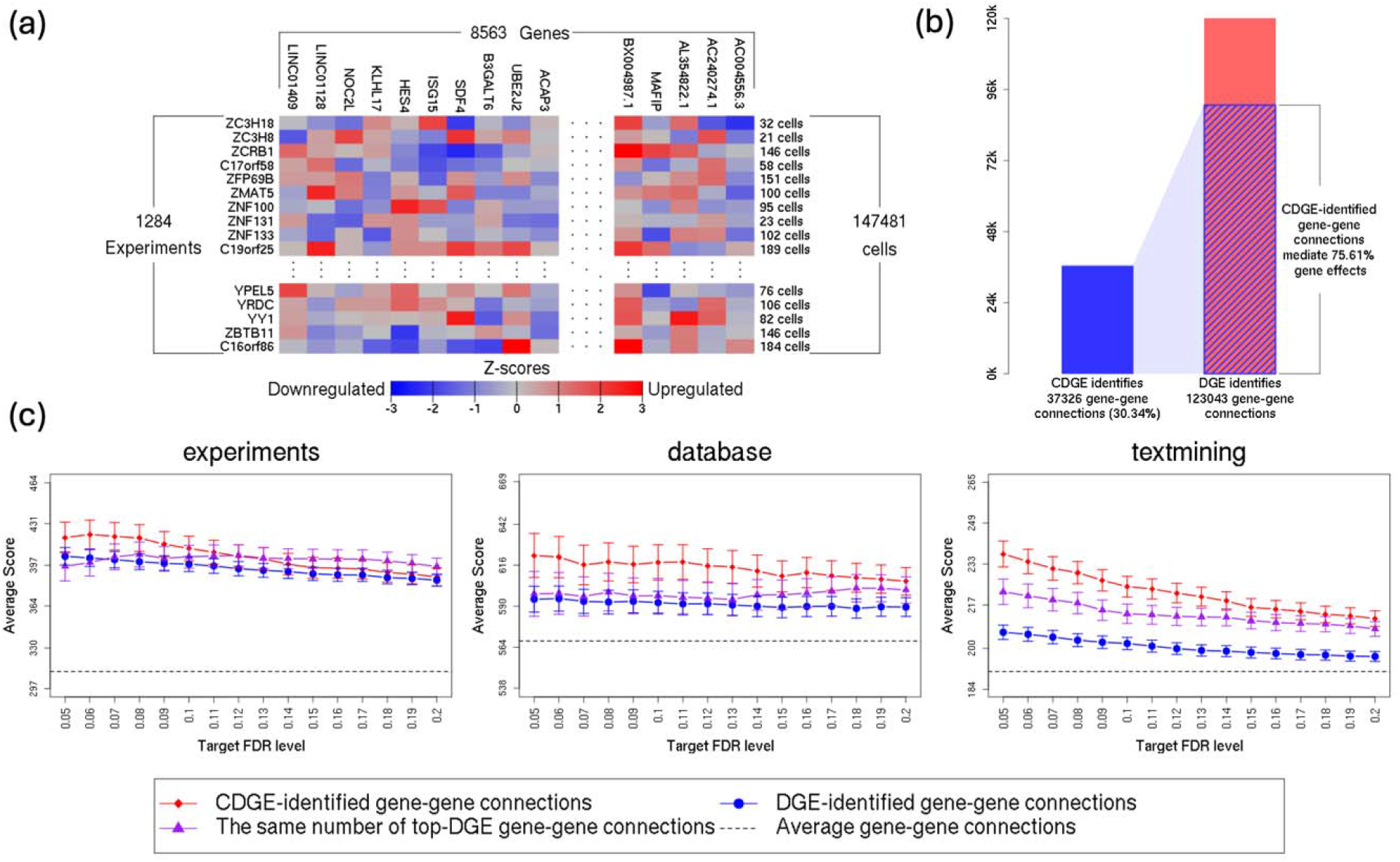
**(a)** Visualization of differential expression Z-scores over 1,284 “P1P2” experiments in comparison to control cells. **(b)** Comparison of the number of CDGE-identified gene-gene connections, the number of DGE-identified gene-gene connections and the number of gene-gene connections whose effect is mediated by CDGE-identified gene-gene connections. **(c)** Confidence scores of protein-protein interactions that correspond to gene-gene connections discovered by knockoff-based CDGE analysis and DGE analysis under different target FDR levels over different evidence channels of the STRING network.

For each experiment, we apply the knockoff-based CDGE analysis on the corresponding Z-scores and expression level correlation matrix **Σ** of all genes to identify gene-gene connections of direct effects. Here, **Σ** is computed via the shrinkage estimation approach (the R package “corpcor”; Schäfer and Strimmer, 2005) using all control cells. We use the same **Σ** throughout all experiments. Under the target FDR level *α* = 0.05, the number of identified gene-gene connections is 37,326 (29.07 per experiments). Details of all these CDGE-identified gene-gene connections are provided in Supplementary Table S3. Such an around 70% decrease is consistent to both the simulation study and analysis of the psoriasis dataset and epilepsy dataset, where most of differentially expressed genes are only of indirect effects or of passenger effects. To validate whether these CDGE-identified gene-gene connections capture direct effects between genes and mediate other gene-gene interactions, we adjust for CDGE-identified genes in each experiment and perform DGE analysis on the remaining genes. Figure 6 (b) shows that these CDGE-identified gene-gene connections only take 30.34% of DGE-identified gene-gene connections in number but explain more than 75%. This suggests that the knockoff-based CDGE analysis can precisely identify gene-gene expressions that capture direct effects and mediate other gene-gene interactions.

We further validate whether the gene-gene connections discovered by the knockoff-based CDGE analysis are more biologically informative. To do so, we perform a series of enrichment analysis with respect to existing biological knowledges. First, we perform an enrichment analysis using the STRING protein-protein interaction (PPI) network (https://string-db.org/, version 12.0; Von Mering et al., 2005) to investigate whether CDGE-identified gene-gene connections are more informative to protein-protein interactions. In the STRING network, 6,857,702 functional associations among 19,699 proteins are annotated in human cells with confidence scores over various evidence channels, including:

1. The *experiments* channel collecting laboratory experiment results imported from BioGRID (Oughtred et al., 2021), DIP (Salwinski et al., 2004), PDB (Berman et al., 2000), IntAct and its partner databases in the IMEx consortium (Orchard et al., 2014). All experiments in these databases are conducted to uncover protein-protein association evidence.
2. Two channels embedding existing knowledge of protein-protein associations, where
  a. The *database* channel records well-established knowledge of protein complexes, pathways and functional connections from KEGG (Kanehisa et al., 2004), Reactome (Gillespie et al., 2022), MetaCyc (Caspi et al., 2020), EBI Complex Portal (Meldal et al., 2022), and Gene Ontology Complexes (Gene Ontology Consortium, 2021).
  b. The *textmining* channel captures the cooccurrence of the gene/protein names in literature collected by the PMC Open Access Subset (up to April 2022), PubMed abstracts (up to August 2022), OMIM (Amberger et al., 2019) and SGD (Cherry et al., 2012).

In the enrichment analysis, we compute the average scores over all channels for protein-protein interactions

- corresponding to CDGE-identified gene-gene connections over *α* = 0.05, 0.06, …, 0.20;
- corresponding to the same number of top-DGE gene-gene connections;
- corresponding to DGE-identified gene-gene connections under FDR levels 0.05, 0.06, …, 0.20

Average scores of all STRING-recorded interactions over all channels are also computed for reference. Figure 6 (c) presents average scores of protein-protein interactions corresponding to different subsets of gene-gene connections in channels *experiments, database* and *textmining*. CDGE-identified gene-gene connections have higher confidence scores than DGE-identified connections under the same target FDR level. This suggests that a lot of biological uninformative gene-gene connections are identified by the DGE analysis, especially in terms of expression patterns and functional connections (in biological pathways and protein complexes). Even if we focus on the most significant DGE-identified connections of the same number as CDGE-identified connections, scores of CDGE-identified connections still dominate in the channels *database* and *textmining*.

We also consider biological pathway libraries under the Molecular Signatures Database (MSigDB; Subramanian et al., 2005; Liberzon et al., 2011; Liberzon et al., 2015) in our enrichment analysis. Specifically, we compare the enrichment of CDGE-identified gene-gene connections to DGE-identified ones in both overlapping and average PPI scores within:

1. All pathways from the Hallmark gene sets (Liberzon et al., 2015);
2. All pathways of malignant cells mined from the Curated Cancer Cell Atlas (3CA) metaprograms (Gavish et al., 2023);
3. All pathways of cancer modules identified in Segal et al. (2004) that changed significantly in various cancer conditions;
4. All pathways in the Biological Process subcollection derived from the Gene Ontology (GO) Consortium (Gene Ontology Consortium, 2021; Gene Ontology Consortium, 2023);
5. All pathways in the Cellular Component subcollection derived from the GO Consortium;
6. All pathways in the Molecular Function Component subcollection derived from the GO Consortium;
7. All pathways in the Human Phenotype Ontology (Köhler et al., 2021).

The number of pathways in each library is presented in the second column of Table 1. In each pathway, we compute the log odd ratio of overlapping for both CDGE-identified gene-gene connections and DGE-identified ones. To compare the degree of enrichment, we perform the one-sided Wilcoxon test (against the null that “CDGE-identified gene-gene connections are at most as enriched as DGE-identified ones”) on log(OR)”s of all pathways in each library and present all the *p*-values in the third column of Table 1. We can find that CDGE-identified gene-gene connections are significantly more enriched than DGE-identified ones in pathways of biological processes, cellular components, molecular functions and human phenotype ontology. In addition, we query PPI scores of these overlapping gene-gene connections in the channels *experiments, database* and *textmining* of STRING, compute the combined score for each gene-gene connection and present one-sided Wilcoxon test *p*-values within different libraries in the fourth column of Table 1. We find that CDGE-identified gene-gene connections have significantly higher PPI scores than DGE-identified ones in pathways of all libraries except the Hallmark gene sets with only 50 pathways. In summary, gene-gene connections identified by CDGE are more biologically informative than DGE.

**Table 1.**
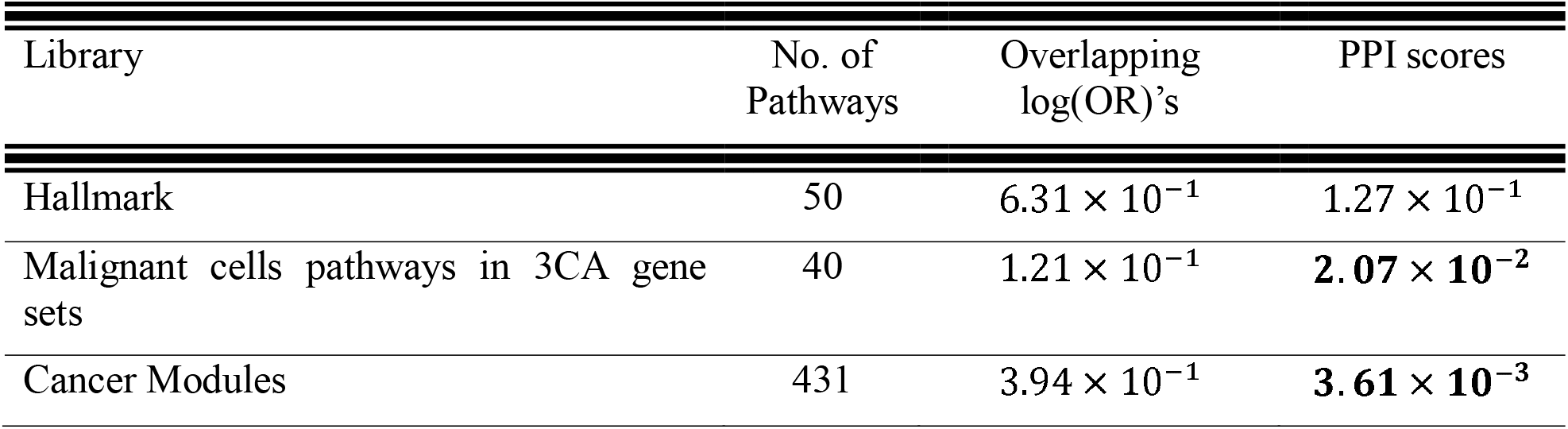
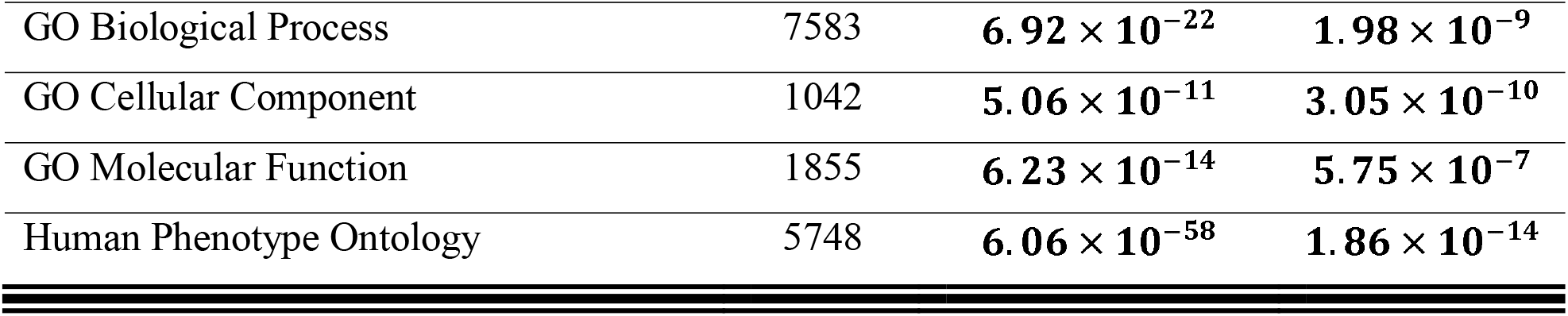
*p*-values of one-sided Wilcoxon tests that compare the degree of enrichment of CDGE-identified gene-gene connections with DGE-identified ones in terms of overlapping log(OR)”s or PPI scores. (Significant results are highlighted in bold)

| Library | No. of Pathways | Overlapping $\log(\text{OR})$’s | PPI scores |
| --- | --- | --- | --- |
| Hallmark | 50 | $6.31 \times 10^{-1}$ | $1.27 \times 10^{-1}$ |
| Malignant cells pathways in 3CA gene sets | 40 | $1.21 \times 10^{-1}$ | **$2.07 \times 10^{-2}$** |
| Cancer Modules | 431 | $3.94 \times 10^{-1}$ | **$3.61 \times 10^{-3}$** |
| GO Biological Process | 7583 | $6.92 \times 10^{-22}$ | $1.98 \times 10^{-9}$ |
| GO Cellular Component | 1042 | $5.06 \times 10^{-11}$ | $3.05 \times 10^{-10}$ |
| GO Molecular Function | 1855 | $6.23 \times 10^{-14}$ | $5.75 \times 10^{-7}$ |
| Human Phenotype Ontology | 5748 | $6.06 \times 10^{-58}$ | $1.86 \times 10^{-14}$ |

## Discussion

In this article, we demonstrate the limitation of differential gene expression (DGE) analysis in learning biological effects of genes. Specifically, we show that differentially expressed genes between cell of different conditions can have either direct effects, indirect effects or passenger effects. This suggests DGE”s lack of ability to distinguish direct effect genes from other differentially expressed genes. To address this issue, we propose the conditional differential gene expression (CDGE) analysis, which infers conditional associations between genes and the trait of interests or among genes with expression level correlations into account. Using the recently developed knockoff approach, we show in extensive simulations that CDGE analysis can identify direct effect genes with guaranteed false discovery rate control while DGE analysis cannot. Analysis of RNA-seq datasets of psoriasis and epilepsy and the experimental perturb-seq dataset validates our statement, as CDGE-identified genes (or gene-gene connections) does mediate effects of other genes (or gene-gene connections) and are more biologically informative.

These findings have direct implications for how RNA-seq results are used in practice. If most differentially expressed genes are passengers rather than drivers, then standard practices are systematically biased toward indirect signals: validating top DEGs experimentally, ranking drug targets by p-value, and interpreting enrichment results across all DEGs all give disproportionate weight to mediated rather than direct effects. The same logic applies to gene-gene regulatory inference from perturb-seq, where the majority of measured downstream changes reflect propagation through the co-expression network rather than direct regulatory effects. CDGE provides a path to redirect follow-up effort toward genes more likely to be biologically causal, with corresponding gains in efficiency for functional validation and therapeutic development. Beyond the specific method, the direct/indirect/passenger/no-effect taxonomy we introduce gives the field a vocabulary for reasoning about what differential expression results mean, regardless of which conditional inference procedure is used to operationalize it.

Our current causal interpretation of CDGE analysis is under the “no unmeasured confounding” assumption. This leaves an open question of how to eliminate confounding effects when using CDGE analysis to investigate biological causality in practice. In RNA sequencing datasets, there usually exist two types of confoundings that blur the identification and interpretation of direct effects, including technical confoundings caused by sequencing depth variation (Sarkar and Stephens, 2021; Su et al., 2023) and biological confoundings caused by unmeasured biological processes. In addition, measurement errors in counting data make inference of direct effects much challenging, because errors not only attenuate direct effects but also mask the conditional independence between the trait of interest and the genes without direct effects (Wang et al., 2018; Zhang et al., 2020). Although out of the scope of this article, it is still desirable to extend CDGE analysis to eliminate unmeasured confounding in distilling causal effects.

Several limitations of the present analysis are worth noting. First, our simulation evaluates CDGE under a single hierarchical generative model, and we have not systematically tested its robustness to correlation-matrix misestimation, model misspecification (for example, when expression follows a negative binomial or zero-inflated distribution), or extreme low *n/p* regimes that arise in some single-cell applications. Practitioners applying CDGE in such regimes should view our FDR guarantees as a target rather than a guaranteed property and verify behavior in their setting where possible. Second, the mediation results we report (particularly the strong claim that adjusting for CDGE-identified genes eliminates the marginal significance of the remaining differentially expressed genes) are based on a single conditioning analysis and have not been benchmarked against random covariate-set baselines; they should be read as suggestive of mediation rather than as a definitive causal decomposition.

Another possible direction lies on the identification of the whole biological pathways by investigating mediated effects. As in Figure 1 and Figure 3 (a), direct effect genes only include the most downstream of biological pathways, while CDGE analysis cannot identify those indirect effect genes on the upstream. With the need to understand the detailed biological mechanisms of how gene effects are mediated and identify drug targets for medical practice with both effectiveness and efficiency, it is vital to develop sequential CDGE analysis with statistical guarantee for biological pathways detection.

## Methods

### Review of existing conditional association analysis

Conditional associations have been extensively studied in literatures of biological data analysis. For example, Ghasemi et al. (2021) used the conditional GCTA (COJO) approach to investigate GWAS signals that depicts conditional associations between genetic variants and free thyroxine (FT4), inflammatory bowel disease (IBD) and human height. Gene regulatory network of conditional associations is also investigated in Li et al. (2021), Ma et al. (2023), Kernfeld et al. (2024) and Tejada-Lapuerta et al. (2025). Specifically, in investigating conditional association between a large set of biological features (denoted by ***X***), and a response of interest (denoted by *Y*) most of literatures impose a multivariate regression model on the conditional model *Y* | ***X***. By doing so, null hypothesis of conditional association (*H*_*C,j*_: *X*_*j*_ ⊥ *Y* | ***X***_−*j*_) is transferred into null hypothesis of corresponding coefficient *β*_*j*_ (*H*_*j*_ : *β*_*j*_ =0) in the regression model, leading various parametric tests and variable selection techniques that infer *X*_*j*_”s contribution to *Y*. Examples include *t*-test, *F*-test, stagewise regression, LASSO (Tibshirani, 1996) and SCAD (Fan and Li, 2001). However, most of inferences based on these methods either lack statistical power or do not rigorously control type-I error rate under high-dimensional settings as in transcriptomics studies with thousands of genes. In addition, they could suffer inference bias when the conditional model *Y* | ***X*** is mis-specified (e.g., true model is nonlinear), which is inevitable due to the lack of domain knowledge.

To overcome the aforementioned limitations, the model-X knockoff filter (Candès et al., 2018) has been developed recently. Imposing no parametric assumption on the conditional model *Y* | ***X***, the model-X knockoff filter only requires knowledge of the distribution of ***X***. Its inference is not only finite-sample valid and powerful but also robust to misspecification of the conditional model *Y* | ***X***. By synthesizing knockoffs 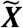 that mimic the dependency structure of ***X*** and do not associate with *Y* beyond ***X***, we use 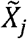 as a control to depict the association between *X*_*j*_ and *Y* if *X*_*j*_ has no direct effect on *Y*(*j* = 1, …, *p*). Thus, the model-X knockoff filter can fairly compare the association test statistics of observed features with their knockoff counterparts and identifies false *H*_*C,j*_”s with rigorous false discovery rate (FDR) control. Based on this novel idea, a series of knockoff methods have been developed. Specifically, as only false *H*_*C,j*_”s correspond to direct effect genes in Figure 1, model-X knockoff filter can prioritize these genes compared to those marginal association methods.

### Knockoff-based CDGE analysis

In this article, our CDGE analysis is conducted based on GhostKnockoff approach with lasso regression, which is proposed in He et al. (2024) and Chen et al. (2024). This allows one to perform CDGE analysis with only summary statistics, defined as Z-scores (or p-values and directions of differential expressions) from usual DGE analysis and the expression level correlations among the genes. The GhostKnockoff approach is originally derived by He et al. (2022) for genome-wide association studies as an analogy of the model-X knockoff filter (Candès et al., 2018). This approach can select substantially contributing genetic variants with valid FDR control in the absence of individual data access. On top of that, Chen et al. (2024) incorporated the BASIL algorithm (Qian et al., 2020) for lasso regression into the GhostKnockoff framework and improved the power. Thus, we adopt the latest GhostKnockoff approach with lasso regression (Chen et al., 2024) to our CDGE analysis for RNA sequencing data. The whole implementation workflow is summarized in Figure 7.

**Figure 7.**
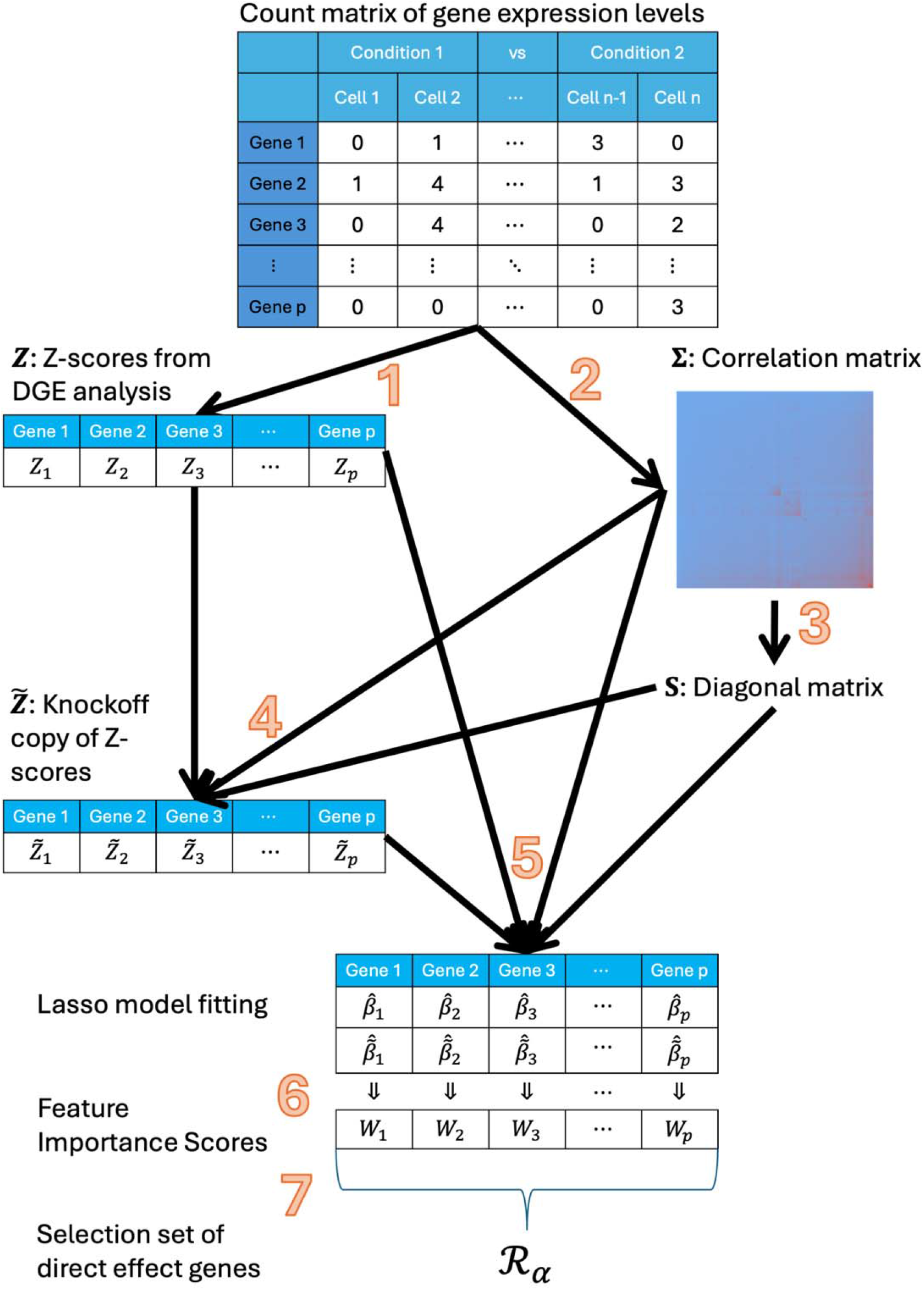
Graphical illustration of the knockoff-based CDGE analysis.

Specifically, as shown in He et al. (2022), Chen et al. (2024) and He et al. (2024), the GhostKnockoff infers the conditional independence hypotheses (*H*_C,*j*_: *X*_*j*_ ⊥ *Y* | ***X***_−*j*_) (*j* = 1, …, *p*) between features *X*_*j*_”s and the response *Y* using two summary statistics:

1. Z-scores ***Z*** = (*Z*_1_, … *Z*_*p*_) that depict the marginal association between each *X*_*j*_ and *Y*;
2. Correlation matrix **∑** that measures the dependency structure among features.

As stated in the “Results” section, genes with direct effects correspond to those false *H*_C,*j*_”s. Thus, we use

1. Z-scores ***Z*** = (*Z*_1_, … *Z*_*p*_)^*T*^ obtained from DGE analysis (e.g., DESeq2, EdgeR and limma) that compares the expression level of each gene across different cell conditions and,
2. correlation matrix **∑** among expression levels of all genes over control cells or all cells, to implement the knockoff-based CDGE analysis and identify direct effect genes. In practice, any existing software that manages to perform DGE analysis can be used to obtain Z-scores (*step 1 in Figure 7*). If the correlation matrix **∑** among gene expression levels is known according to domain knowledge or reference studies, we can directly use it as in our simulated experiment. When such existing knowledge is not available, the correlation matrix **∑** can be estimated either via the factor model as in the analysis of psoriasis and epilepsy datasets or via the shrinkage estimation approach provided in the R package “corpcor” (Schäfer and Strimmer, 2005) as in the analysis of the experimental perturb-seq dataset (*step 2 in Figure 7*).

With Z-scores ***Z*** and correlation matrix **∑**, the next step is to compute a diagonal matrix ***S*** = diag((*s*_1_, … *s*_*p*_) for knockoffs generation under a particular construction regime. Possible regimes include minimizing the mean absolute correlation (Barber and Candès, 2015; Candès et al., 2018), maximizing the entropy (Gimenez and Zou, 2019) and minimum variance-based reconstructability (Spector and Janson, 2022). In practice, we compute the matrix ***S*** under the maximizing the entropy criterion, although other criteria are also allowed (*step 3 in Figure 7*).

We then synthesize the knockoff copy 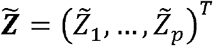 of Z-scores following He et al. (2022) that

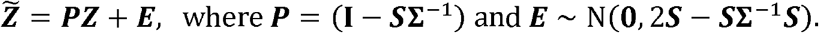

Here, 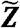 serve as synthetic controls where 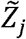 captures all indirect effects from the *j*-th gene (*X*_*j*_) to the response *Y* through other genes. Thus, for those genes without direct effects (whose *H*_C, *j*_’s are true),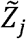”s will be similar to *Z*_*j*_’s; for those genes with direct effects (whose *H*_C,*j*_’s are false), *Z*_*j*_’s will dominate 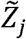”s by capturing direct effects.

Although it is possible to select direct effect genes with FDR control by directly comparing *Z*_*j*_”s and 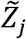”s (He et al., 2022), doing so may suffer power loss due to noises brought by indirect effects. To circumvent this, we adopt the strategy in He et al. (2024) and Chen et al. (2024) to fit a lasso regression model and obtain estimates 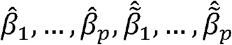, which depict the direct effects of genes and their knockoff copies to the disease, trait of interest or the perturbed gene. Specifically, with 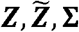, the lasso estimates 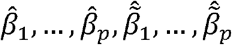 are obtained by solving the optimization problem,

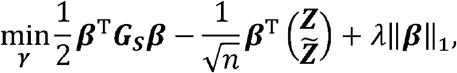

where 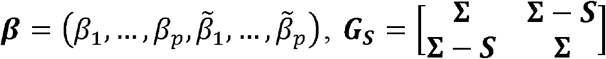 and *λ* is computed via the lasso-min method of Chen et al. (2024) (*step 5 in Figure 7*).

As 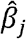 captures the direct effect of the *j*-th gene while 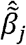 capture the direct effect of the synthetic control that mimics the *j*-th gene, we have

- for genes without direct effects,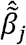”s will be similar to 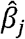”s;
- for genes with direct effects, 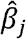 “s will dominate 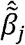”s in magnitude.

Thus, we calculate the feature importance score as

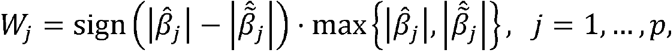

such that *W*_*j*_”s tend to be large and positive for direct effect genes and *W*_*j*_”s fluctuate around 0 otherwise (*step 6 in Figure 7*).

Finally, we apply the knockoff filter (Barber and Candès, 2015; Candès et al., 2018) to select direct effect genes under the target FDR level *α* as

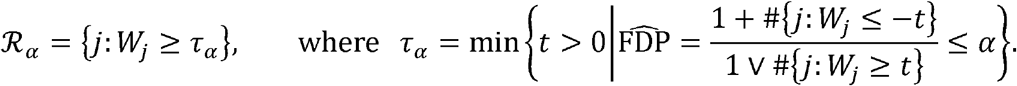

Theoretical studies in Chen et al. (2024) have shown that the selection set of direct effect genes ℛ _*α*_ has valid control of FDR (*step 7 in Figure 7*).

In the accompanying R package “KnockoffCDGE” for implementing knockoff-based CDGE analysis, the output of the R function “KnockoffCDGE” is q-values of all features,

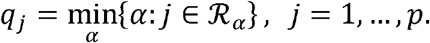

Here, *q*_*j*_ characterizes the minimum target FDR level at which the *j*-th gene is identified and thus users can obtain ℛ _*α*_ = {*j* : *q*_*j*_ ≤ α}

## Experiments

### Data generation mechanism in simulated experiments

For each replicate, we generate the expression levels of *p* = 500 genes in *n* = 5, 000 cells as follows.

1. Generate expression levels of indirect effect genes as *X*_indirect,1_ = (*X*_1_, … *X*_20_)^T^ where *X*_1_, … *X*_20_ are i.i.d. random variables from Poisson distribution *Pois*(1).
2. Generate expression levels of no effect genes as *X*_null, 1_ = (*X*_21,_ … *X*_80_)^T^ = (*C*_null, 1_ + *E*_21,_ … *C*_null, 1_+ *E*_80_)^T^ where *E*_21,_ … *E*_80_ ∼ *Pois* (1 − *ρ*_1_) and *C*_null, 1_ ∼ *Pois* (1 − *ρ*_1_) are mutually independent with *ρ*_1_ = 0.7.
3. Generate expression levels of passenger genes as *X*_passenger,1_= (*X*_81,_ … *X*_90_)^T^ from

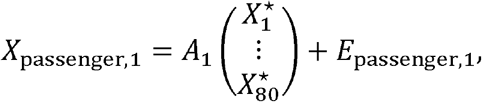

where *E*_passenger,1_ is a 10-dimensional vector of i.i.d. *Pois*(1) random variables, *A*_1_ is a random matrix whose entries are i.i.d. samples from Bernounli distribution *Bern* (0.2), 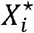 “s are independent random variables from *B* (*X*_*i*,_ *q*_*i*_) with *q*_*i*_ ∼ *Unif* [0, 1/4] for *i* = 1, …, 80.
4. For *l* = 2, … 5,
  a. Generate expression levels of indirect effect genes as *X*_indirect,*l*_ = (*X*_1+90 · (*l*−1),_ … *X*_20+90 · (*l*−1)_^T^ from

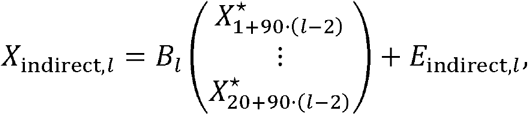

where *E*_indirect,*l*_ is a 20-dimensional vector of i.i.d. *Pois*(1) random variables, *B*_*l*_ is a random matrix whose entries are i.i.d. samples from Bernounli distribution 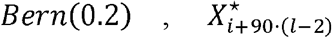 “s are independent random variables from *B* (*X*_*i*+90 · (*l*−2),_, *q*_*i*+90 · (*l*−2)_ with *q*_*i*+90 · (*l*−2)_ ∼ *Unif* [ 0,1/4] for *i* = 1, …, 20.
  b. Generate expression levels of no effect genes as *X*_null_, *l* (*X*_21+90 · (*l*−1),_, … *X*_80+90 · (*l*−1)_^T^= (*C*_null, *l*_ +*E*_21+90 · (*l*−1),_ …, *C*_null, *l*_ + *E*_80+90 · (*l*−1)_^T^ where *E*_21+90 · (*l*−1)_ …, *E*_80+90 · (*l*−1)_ ∼ *Pois*(1 − ρ_*l*_) are mutually independent with 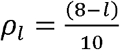.
  c. Generate expression levels of passenger genes as *X*_passenger, *l*_ = (*X*_81+90 · (*l*−1),_ … *X*_90+90 · (*l*−1)_^T^ from

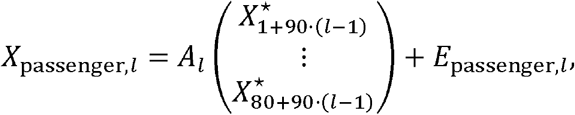

where *E*_passenger,1_ is a 10-dimensional vector of i.i.d. *Pois*(1) random variables, *A*_*l*_ is a random matrix whose entries are i.i.d. samples from Bernounli distribution 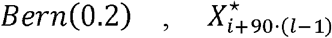 “s are independent random variables from *B*(*X*_*i*+90 · (*l*−1),_ *q*_*i*+90 · (*l*−1)_ with *q*_*i*+90 · (*l*−1)_ ∼ *Unif* [ 0,1/4] for *i* = 1, …, 80.
5. Generate expression levels of direct effect genes as *X* _direct_ = (*X*_451,_ … *X*_500_)^T^ from

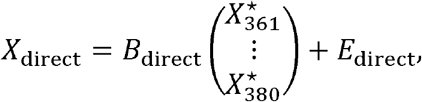

where *E*_direct_ is a 50-dimensional vector of i.i.d. *Pois*(1) random variables, *B*_direct_ is a random matrix whose entries are i.i.d. samples from Bernounli distribution 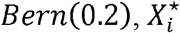 “s are independent random variables from *B* (*X*_*i*,_ *q*_*i*_) with *q*_*i*_ ∼ *Unif* [ 0,1/4] for *i* = 361, …, 380.

### Additional experiment details of the epilepsy dataset

In this section, we provide details of the mice experiment that collects the epilepsy dataset. This experiment is performed through dissection of 4 heterozygous (Scn8a+/-) mice and 4 wild-type (Scn8a+/+) mice as controls (RRID: IMSR_JAX:003798). All mice are 56-57 days old when their brains are extracted, where the cuts are positioned at the bregma coordinates 1.045mm to - 1.055mm and subcortical regions below the corpus callosum are removed. The extracted tissues are then stored in MACS Tissue Storage Solution with 45 *μ* M Actinomycin D. After isolating nuclei using MACS sorting, we fix brain nuclei of mice using Parse biosciences fixation kit, run them thought the Parse biosciences WT barcoding kit (capacity: 100k cell/nuclei). We then sequence cDNA of different brain cells under Seqmatic Novoseq X Plus Run, which returns Fastq files. For statistical inference in R, we finally pass the Fastq files through Parse Pipeline V1.1.0 to demultiplex the raw reads into a RDS file.

## Code Availability

To facilitate public access to knockoff-based CDGE analysis, we have developed the R package “KnockoffCDGE” to provide an integrated implementation, which is freely available at https://github.com/GuJQ5/KnockoffCDGE.

## Acknowledgments

The study was supported by NIH/NIA award AG089509, AG066206 and AG066515, NIH-NINDS K08 award NS119800, Stanford MCHRI Tashia and John Morgridge Endowed Faculty Scholar Award and McKnight Foundation Award.

